# A Codon Constrained Method for Both Eliminating and Creating Intragenic Bacterial Promoters

**DOI:** 10.1101/2021.08.04.454459

**Authors:** Dominic Y. Logel, Ellina Trofimova, Paul R. Jaschke

## Abstract

Future applications of synthetic biology will require refactored genetic sequences devoid of internal regulatory elements within coding sequences. These regulatory elements include cryptic and intragenic promoters which may constitute up to a third of predicted *Escherichia coli* promoters. Promoter activity is dependent on the structural interaction of core bases with a σ factor. Rational engineering can be used to alter key promoter element nucleotides interacting with σ factors and eliminate downstream transcriptional activity. In this paper, we present <u>CO</u>don <u>R</u>estrained <u>P</u>romoter <u>S</u>il<u>E</u>ncing (CORPSE), a system for removing intragenic promoters. CORPSE exploits the DNA-σ factor structural relationship to disrupt σ^70^ promoters embedded within gene coding sequences, with a minimum of synonymous codon changes. Additionally, we present an inverted CORPSE system, iCORPSE, which can create highly active promoters within a gene sequence while not perturbing the function of the modified gene.

## Introduction

Building predictable and orthogonal transcriptional networks is a central goal in synthetic biology. The mass re-engineering of complex native genetic sequences into predictable and rational systems, through a process known as refactoring, is dependent on maintaining biological function while removing cryptic regulation and genetic overlaps ^1, 2^. Refactoring is a broad term in synthetic biology applying to multiple approaches which simplify genetic systems on a multi-gene or genomic level. Many examples of refactoring have been demonstrated in both multi-gene pathways ^3–5^, and bacteriophage ^6–8^.

The ongoing design of synthetic and modified genetic systems is complicated by internal genetic regulation, such as promoter sequences internal to a coding sequence, which can create deviations from predicted outcomes ^3, 9, 10^. While the majority of bacterial promoters are in intergenic regions, many promoters are intragenic, overlapping an upstream coding sequence. Well-known examples of intragenic promoters include the *micLp* promoter within the *Escherichia coli cutC* gene driving *micL* small RNA transcription ^11^ and the *trpCp* promoter within *trpD* driving *trpC* transcription ^12^.

Recent advances in methods to detect transcription start sites (TSS) across the genome has suggested that *E. coli* transcription under multiple conditions contains nearly 15,000 TSS ^13^ with 32 % located internal to coding sequences. It is currently unclear how biologically relevant these newly discovered promoters are, and recent work with other experimental methods has shown far fewer promoters in *E. coli* ^14^.

Bacterial promoter sequences are typically made of two core sequence elements, the −35 and −10 elements. The core elements bind to the primary specification component within the RNA polymerase (RNAP) holoenzyme, the σ factor. Transcription is initiated when a σ factor binds to dsDNA at the −35 element to promote DNA isomerisation to ssDNA at the −10 element which drives transcription events. Promoter DNA-protein interactions are not limited to the two core elements as holoenzyme interactions can be enhanced or replaced through accessory sequences such as the extended −10 TGn motif and UP elements, which interact with the bacterial σ factor and α subunits respectively ^15^. Promoter elements interacting with the housekeeping σ^70^ are the most well-characterised ^16^, and have been demonstrated to bind specifically with their target promoters through DNA-protein interactions such as hydrogen bonding, Van der Waals forces, and stacked cation-π bonds ^17, 18^. The −35 element nucleotide sequence identity is a relatively promiscuous interaction because the σ factor mainly recognises the −35 element’s nucleotide backbone ^18^, while in comparison the −10 element DNA-protein interaction is more discriminatory. The −10 element DNA-protein interaction relies on two bases in the sequence, ^−11^A and ^−7^T, which base-flip and change orientation from the DNA sequence stack to interact within two highly specific binding pockets in the σ factor ^17^. The specific nature of DNA-σ factor interactions leaves opens a rational approach to silently eliminate intragenic promoter sequences within a coding sequence by eliminating DNA-σ interactions through synonymous mutations.

Typically, intragenic promoters have been eliminated during refactoring by extensive randomization of synonymous codons of overlapping coding sequences ^5^. However, the mass randomisation of codons in a coding sequence can have unpredictable deleterious effects when specific codons play a play a critical role in translation rate ^19^, or protein folding and function ^20–23^. Furthermore, synonymous codon changes have been seen to create mRNA toxicity independent of translation ^24^. Therefore, a rational approach that minimizes codon changes while still effectively disrupting internal promoters would be highly desirable.

In this work, we present a codon-aware promoter removal method called <u>CO</u>don <u>R</u>estrained <u>Pr</u>omoter <u>S</u>il<u>E</u>ncing, or CORPSE. CORPSE aims to disrupt internal promoters through minimal synonymous codon changes. We show the potential of the CORPSE method through erasing the *trpCp* intragenic promoter without perturbing the overlapping *trpD* gene function in *E. coli*. We also present inverted CORPSE (iCORPSE), which is capable of silently inserting a promoter within a coding sequence with only a few synonymous codon modifications. We demonstrate iCORPSE by creating a new promoter within the fluorescent reporter gene mCherry without disrupting its function. Together, these two methods present a platform for precisely eliminating intragenic promoter elements from coding regions during refactoring, and the construction of new intragenic promoters for next-generation compressed genetic circuit design.

## Results and Discussion

### The CORPSE method can eliminate promoter activity using only synonymous mutations

To create the CORPSE method, we first gathered σ^70^ promoter sequence data from the *E. coli* K-12 database RegulonDB ^25^. From this list of promoters we removed all promoter elements varying from the standard hexameric nucleotide length. The remaining promoters were used to create a position specific scoring matrix (PSSM). We used the PSSM to score promoters based on how close they conformed to the σ^70^ consensus.

Next, we created an algorithm to consider the hexameric sequences in a promoter as codons, as would be the case for intragenic promoters, and then to generate alternative sequences that are limited to only synonymous codon variants at each position.

To test the CORPSE algorithm’s ability to reduce or eliminate a σ^70^ promoter, we chose the intragenic promoter *pB* within the bacteriophage ϕX174 genes A/A* as a model ^26^. We used the CORPSE algorithm to generate a list of 11 possible alternative sequences each for the −35 and −10 promoter elements that would be expected to reduce promoter activity while maintaining synonymous codons of overlapping coding sequences A/A* (Figure 1A and 1B). The −35 element synonymous codon PSSM scores ranged from −0.43 to 4.36 which are in the 25^th^ - 70^th^ percentiles of all *E. coli* σ^70^ promoters (Figure S1A), while the unmodified sequence score (5.36) is in the 89^th^ percentile. The −10 element PSSM scores ranged from −0.68 to 6.34, placing them in the 10^th^ - 70^th^ percentiles of all *E. coli* promoters, while the unmodified sequence score (4.44) was in the 40^th^ percentile (Figure S1B). We generated promoter elements using CORPSE that reduced the score to less than 2 (33^rd^ percentile) for the −35 and less than 1 (18^th^ percentile) for the −10 element, respectively (Figure 1C). We combined these selected elements combinatorically to generate 16 variants to examine further (Table S1).

**Figure 1:**
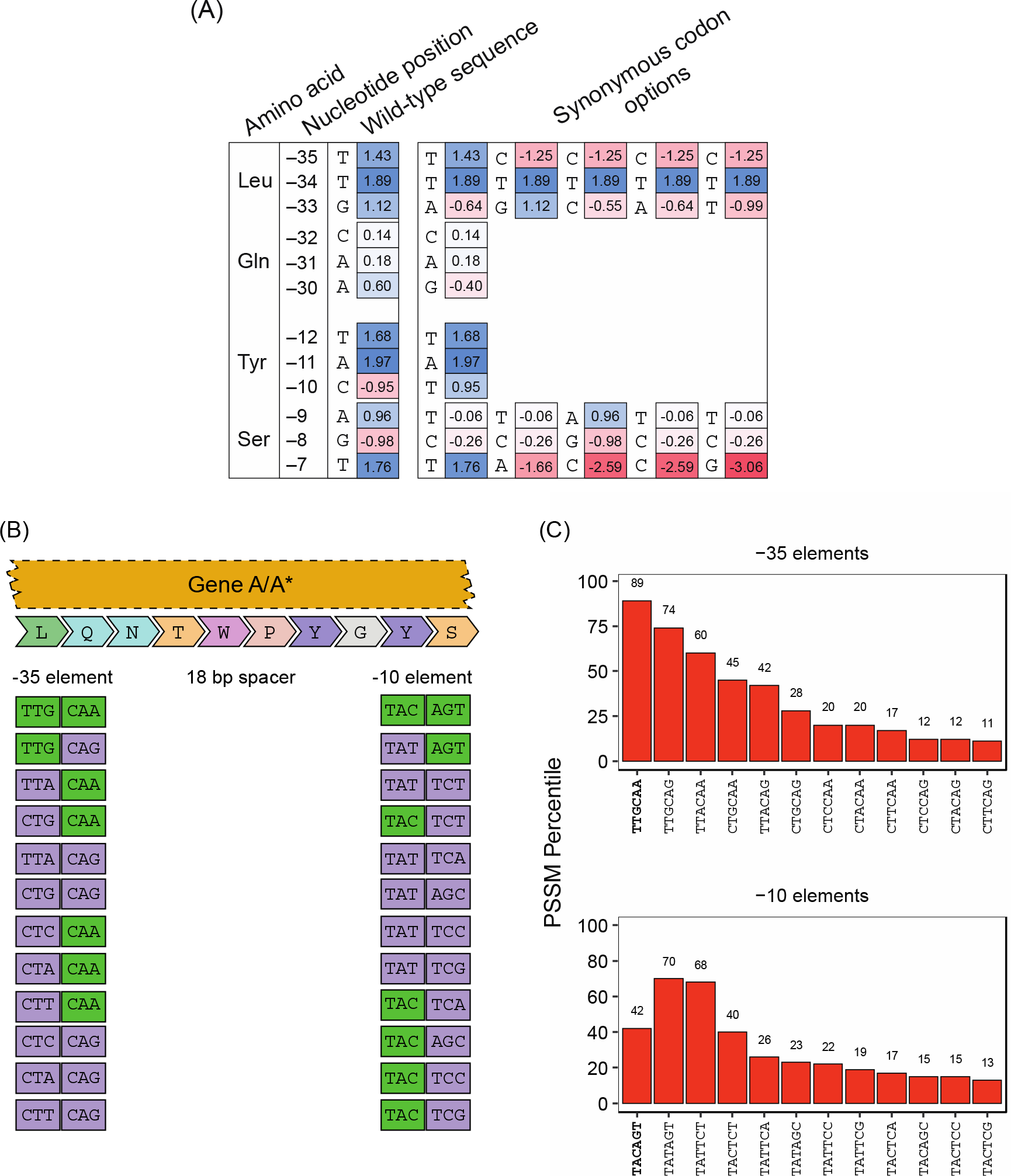
CORPSE algorithm. (A) List of all possible CORPSE modifications for *pB* with their PSSM score noted. Colors denote calculated score within each position in PSSM, high to low scores are shown as blue to red gradient. (B) List of all selected CORPSE variants used to eliminate *pB* activity with wild-type codons shown in green and variants in purple. Codons shown corresponded to translated *pB* sequence within ϕX174 genes A/A* (C) PSSM percentile for each possible variant with percentile shown above each sequence. The wild-type sequences are shown in bold.

To determine if CORPSE mutations reduced the activity of the *pB* promoter, the 16 variants were assembled into the pJ804 plasmid upstream of superfolder GFP (sfGFP). The plasmid also contained mCherry in the reverse direction to sfGFP under control of a constitutive promoter (Figure 2A) for expression capacity normalization ^27^.

**Figure 2:**
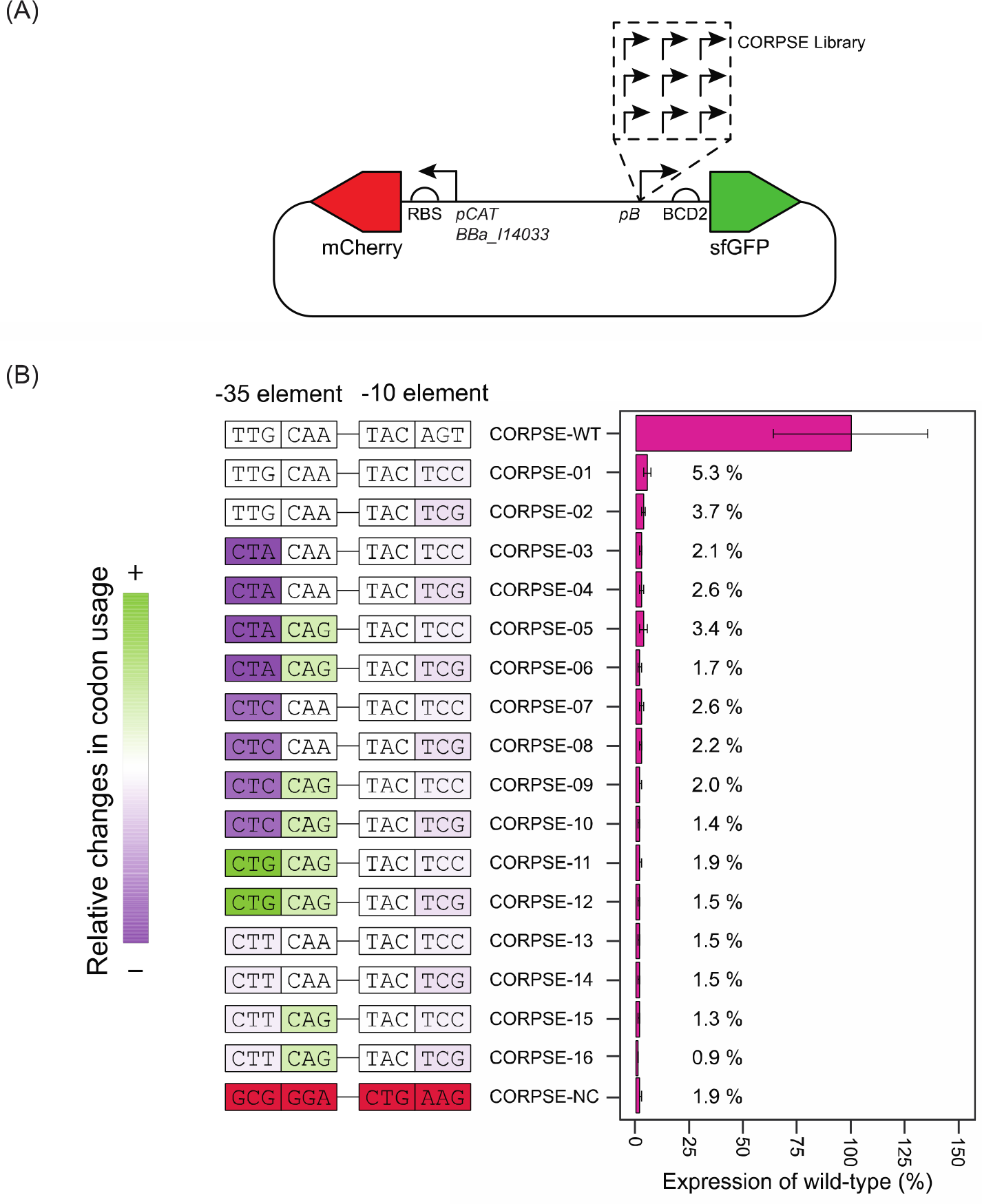
Low scoring promoter variants dramatically reduce *pB* promoter activity, while only making synonymous changes to overlapping codons. (A) CORPSE reporter plasmid contains constitutively expression mCherry as an internal control while wild-type and CORPSE mutants of *pB* driven variable sfGFP expression. (B) All mutations to *pB* reduced activity of the promoter within the reporter plasmid with the minimal changes to the −10 element in CORPSE-02 displaying only 5.3 % of wild-type activity. Broader modifications to the sequence changing both promoter elements further decreased expression with normalized expression being reduced to 0.9 % of wild-type. CORPSE-NC represents a negative control with selected nucleotides being worse scoring choices at each position, ignoring codon restraints. Normalised sfGFP compared to wild-type is shown as %.

Plasmids were built, transformed into *E. coli*, and their fluorescence measured using flow cytometry. The median sfGFP values for each variant were normalized to mCherry and compared to the wild-type construct. In all variants, sfGFP expression was significantly reduced to 0.9 % −5.3 % of wild-type (p values < 0.005) (Figure 2B). The CORPSE alterations to *pB* were highly effective as the minimal mutations within CORPSE-01, a ^−9^AGT^−7^ to ^−9^TCC^−7^ modification, were disruptive enough to remove 95 % of the promoter’s function. These results were not unexpected as the two most conserved sequences in the −10 element, ^−11^A and ^−7^T, are found 88.5 % and 80.2 % of the time respectively in σ^70^ promoters ^25^. These two nucleotides base flip out of the DNA stack during isomerisation and bind with unique pockets within the σ factor.

The likely driver behind the reduced expression was the altered −7 base which canonically binds within an hydrophilic pocket in the σ factor which disallows pyrimidine nucleotides binding and structurally excludes ^−7^C bases, a mutation which was present in half of the CORPSE variants (Figure 2B) ^17^. While the −11 base is a useful target for rational engineering due to its binding constraints ^28^, the synonymous codons for tyrosine (TAT or TAC) within the gene A/A* sequence overlapping the promoter only allowed the third nucleotide to vary, and changing the C to a T would have increased, not decreased promoter strength.

While the −7 modifications likely drove expression changes, the non-base flipping nucleotides, −10 to −8, still bind to the σ factor through sugar-phosphate backbone interactions, as well as potential ^−8^A nucleotide van der Waal force contact with the σ factor ^17^. These structural interactions bring potential for further reductions in ideal DNA-σ factor interactions, however, are likely limited as they interact via their backbone, not nucleotide structure.

### CORPSE can silently eliminate *trpCp* intragenic promoter without affecting overlapping *trpD* gene function

To determine if this method could be more broadly applied to *E. coli* refactoring, and to test if the codon changes affect the overlapping gene function, we applied the CORPSE method to the *trpCp* promoter within the tryptophan biosynthesis operon (Figure 3A). The *trpCp* promoter is located within the *trpD* gene sequence ^12^ and provides additional transcriptional current to drive basal expression of the trpCBA genes ^12, 29^. The wild-type sequence of *trpCp* has a −35 and −10 PSSM score of 3.2 and 1.6 respectively which placed them within the 32^nd^ and 21^st^ percentiles (Figure S1A and S1B), corresponded with the known low strength of *trpCp* ^12, 29^. We applied the CORPSE method to generate a set of all possible variants that would significantly reduce the promoter strength while retaining the same amino acids of the overlapping TrpD protein.

**Figure 3.**
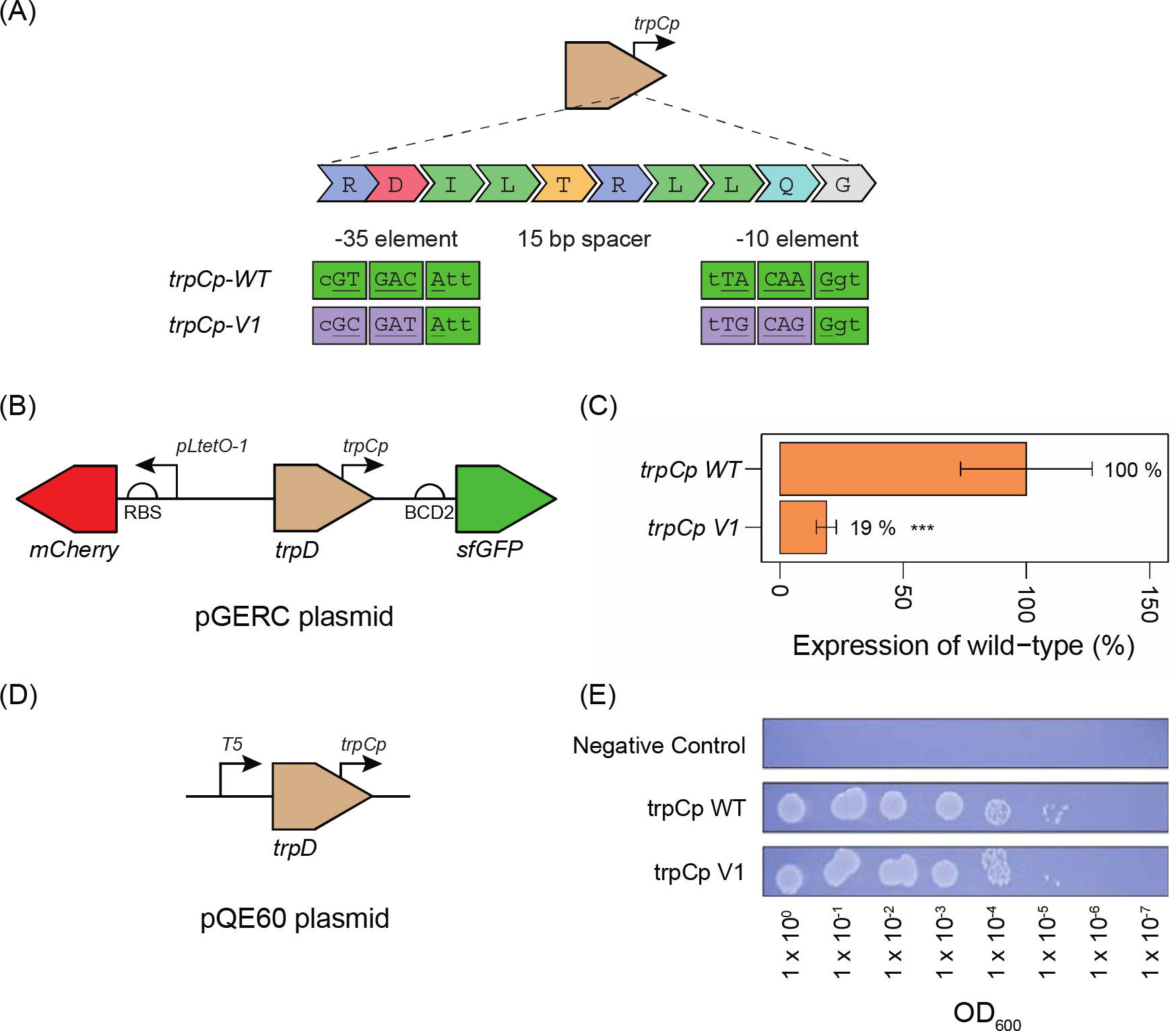
Wild-type and CORPSE variant supplementation of *trpCp* containing *trpD* constructs reduced reporter expression and maintained gene functionality. (A) The *trpCp* promoter within *trpD* was modified through CORPSE to eliminate the promoter. Wild-type codons shown in green and synonymous mutations in purple. Promoter hexamers underlined within codons. (B) Wild-type and CORPSE V1 *trpD* genes were inserted upstream of *sfGFP* in a pGERC plasmid to drive reporter expression. (C) CORPSE modification of *trpCp* successfully reduced reporter expression by 81 % when normalized to internal mCherry expression with p values < 0.005 shown as ***. (D) Wild-type and CORPSE V1 *trpD* gene were inserted into qQE-60 plasmids and expressed in the Δ*trpD* KEIO collection strain JW1255-1 to assess gene function effects of the modification. (E) Overnight cultures of empty pQE60 and the two *trpD* constructs were grown on tryptophan deficient M9 media after normalization and serial dilution. No phenotypic effects were detected from the *trpCp* modifications as no growth defects were apparent.

We selected one variant which reduced the −35 and −10 scores to the 1^st^ and 4^th^ percentiles respectively for further testing. We assembled the wild-type (pGERC::*trpCp-WT*) and variant *trpD* (pGERC::*trpCp-V1*) gene sequences into the pGERC plasmid upstream of a sfGFP gene to probe the expression effects of our modifications (Figure 3B). Again, an mCherry gene under constitutive promoter was located in the opposite orientation on the plasmid as expression normalization control. The two strains were analysed using flow cytometry and the results showed pGERC::*trpCp-V1* displayed an 81 % reduction in sfGFP fluorescence compared to pGERC::*trpCp-WT* (Figure 3C). As with the *pB* variants, the pGERC::*trpCp-V1* variant contained minimal changes to the wild-type sequence with only two nucleotide modifications in the −35 element and three modifications in the −10 element.

To test whether the synonymous mutations made to disrupt the *trpCp* promoter had any deleterious effects on the overlapping *trpD* gene function, we assembled the wild-type and modified *trpD* genes into a pQE60 plasmid (Figure 3D) and attempted to complement a strain from the KEIO collection with a *trpD* disruption (JW1255-1). The two complemented strains, along with an empty pQE60 plasmid control, were grown overnight in LB media followed by plating on M9 minimal media lacking tryptophan. After overnight growth we observed identical growth patterns for strains pQE60::*trpCp-WT*) and (pQE60::*trpCp-V1*) on both LB and tryptophan deficient M9 minimal media (Figure 3E) demonstrating that modified *trpD* was functionally equivalent to wild-type *trpD* under these conditions. While both complemented strains grew on M9 minimal medium lacking tryptophan, the strain containing the empty plasmid control did not grow.

While wild-type *trpCp* already contained a non-optimal nucleotide in the −7 position, ^−7^G, the promoter contained a consensus ^−11^T. We were able to break this consensus sequence by altering ^−13^TTA^−11^ to ^−13^TTG^−11^ in the −10 element. The nucleotide with the largest predicted effect was the alteration of ^−37^AGT^−35^ codon to ^−37^AGC^−35^. This in theory affected the interactions of the σ subunit to the nucleotide and backbone atoms at the −36 and −35 positions ^18^. In addition to significantly reducing sfGFP expression, when the wild-type and variant *trpD* genes were expressed in a Δ*trpD* cell strain, no changes in cell growth were detected when grown on minimal media. This suggests that the specific synonymous mutations used had no effect on *trpD* functionality.

### Inverted CORPSE (iCORPSE) can silently create new promoters within gene sequences

We next tested the feasibility of inverting CORPSE to create a functional promoter within a coding sequence without disturbing the function of the overlapping gene. We first used the promoter prediction software BPROM ^30^ to analyse the complementary strand of the mCherry gene to identify any sequences already displaying promoter-like characteristics that could be modified to generate a *de novo* promoter directing transcription in the opposite direction to mCherry and its promoter. Two promoters were predicted through that software, designated PROM-1 and PROM-2 (Figure 4A). In our experience the BPROM software often predicts sites that are not actually promoters *in vivo* in addition to identifying sites that are true promoters^26^. We also manually identified a site (PROM-3) with two hexameric sequences separated by 17 bp that could be synonymously modified towards the σ^70^ consensus sequence (Figure 4A and B, and Table 1). Analysis of the proto-promoters showed they had a range of PSSM scores that put the −35 elements in the 10^th^ – 35^th^ percentiles and the −10 elements in the 4^th^ – 95^th^ percentiles of *E. coli* σ^70^ promoters (Table 1).

**Fig. 4:**
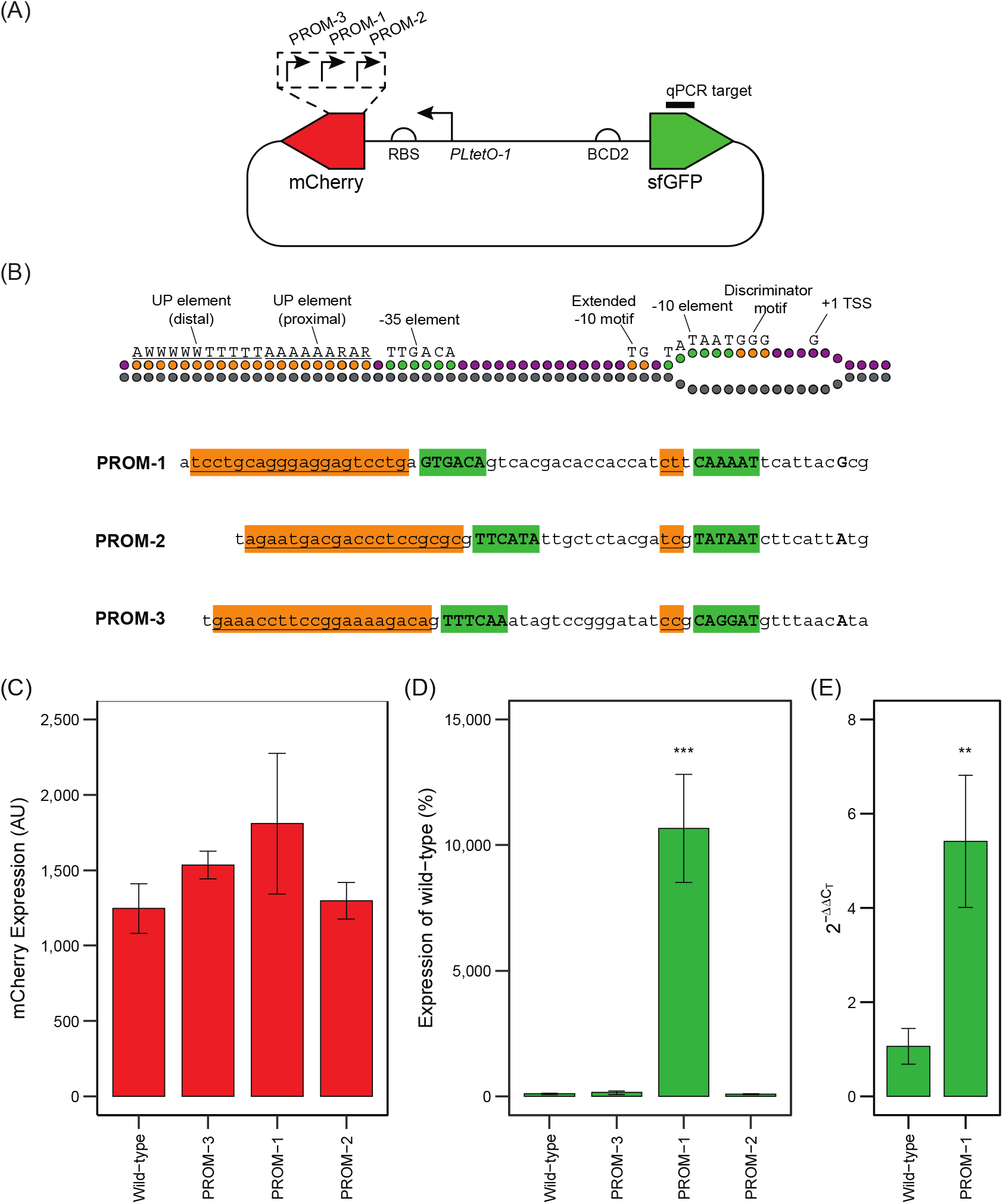
iCORPSE driven sfGFP expression. (A) Reporter plasmid for iCORPSE has constitutively driven mCherry expression and variable sfGFP expression driven by synonymous mutations within the mCherry antisense reading frames. Three potential promoters were engineered into mCherry to drive sfGFP expression. The qPCR target for sfGFP is shown. (B) Engineered iCORPSE sequence aligned against structural map of σ^70^ promoter sequences. Core binding elements shown in green and accessory binding elements shown in yellow. (C) Fluorescence from sfGFP expression across mCherry variants compared to wild-type mCherry sequence. (D) Mean mCherry expression and standard deviation. (E) Log2 fold-change results for sfGFP expression normalized against housekeeping cysG. P values < 0.01 and <0.001 are shown as ** and ***, respectively.

**Table 1:** iCORPSE sequences.

| Promoter variant | -35 element | -10 element | PSSM score |  |  |  |  |
| --- | --- | --- | --- | --- | --- | --- | --- |
|  |  |  | -35 | Percentile | -10 | Percentile | combined |
| PROM-1 | GTGACG (wt) | CAAAAT (wt) | 2.24 | 37<sup>th</sup> | 4.07 | 39<sup>th</sup> | 6.31 |
|  | GTGACA | CAAAAT | 3.24 | 54<sup>th</sup> | 4.07 | 39<sup>th</sup> | 7.31 |
| PROM-2 | CTCATA (wt) | TATAAT (wt) | 1.27 | 24<sup>th</sup> | 8.35 | 95<sup>th</sup> | 9.62 |
|  | TTCATA | TATAAT | 3.95 | 65<sup>th</sup> | 8.35 | 95<sup>th</sup> | 12.3 |
| PROM-3 | CTTCAG (wt) | CTGGGT (wt) | -0.43 | 10<sup>th</sup> | -7.37 | 4<sup>th</sup> | -7.80 |
|  | TTTCAA | CAGGAT | 3.25 | 54<sup>th</sup> | 1.30 | 19<sup>th</sup> | 4.55 |

Next, we used the CORPSE algorithm to generate a list of synonymous codon changes increasing the cumulative PSSM score across both elements, followed by modifying the mCherry sequence within the pGERC plasmid ^31^ (Figure 4A) to reflect the recoded PROM-1, PROM-2, and PROM-3 sequences. Measurements of mCherry and sfGFP fluorescence from cells carrying the plasmids showed PROM-2 and PROM-3 modifications resulted in no detectable change to sfGFP expression compared to the control plasmid (Figure 4D). In contrast, PROM-1 dramatically increased sfGFP expression to 10,664 % over the control plasmid (Figure 4D). The synonymous codon changes used to create the promoter had no significant (p value ≤ 0.01) effect on mCherry fluorescence (Figure 4C). To confirm the increased sfGFP expression was due to increased transcription of the sfGFP gene we extracted RNA and performed qPCR with a primer set targeted to the sfGFP coding sequence (Figure 4E). RNA abundance of the sfGFP transcript was found to be 5.4-fold increased when the modified PROM-1 sequence was present in mCherry versus the wild-type mCherry sequence (Figure 4E).

We cannot fully explain the differences in sfGFP expression from the different iCORPSE variants using the CORPSE scoring scheme. All iCORPSE sequences contain the crucial −10 element nucleotides ^−11^A and ^−7^T bases (Figure 4B). In addition, no iCORPSE sequence displays the canonical A/T tract repeats of an UP region ^32^ or has a canonical extended −10 TGn motif ^33^ and therefore has not clear enhancer motifs. The difference in spacer length may explain the differential effects in promoter activity because BPROM-1 contains 19 bp, BPROM-2 contains 13 bp, and PROM-3 contains a 17 bp spacer. Through a spacer sequence explanation, the extended BPROM-1 spacer could permit more dynamism in RNAP binding which was more restricted in the other two iCORPSE variants ^34^.

Most attempts to predict strength from sequence identities have utilised statistical methods with mixed success ^35–37^. The overall biological complexity of the system complicates prediction which in the future may be eliminated by large measurement sets and machine learning approaches that are only now becoming available ^38–41^.

A future direction for synthetic genomics research will be to progress from ordered genome architectures where all regulatory elements and coding domains are separate from each other, and towards compressed architectures with overlapping coding and regulatory sequences. The advantages of a compressed architecture would be reducing overall genome size ^42^, and translationally couple gene expression ^43, 44^ permitting greater genetic content within a reduced length. Intragenic overlapping promoters could potentially shield them from mutation but any changes created with synonymous changes could be easily undone by synonymous mutation.

In conclusion, in this study we used a rational system to eliminate function from a promoter internal to a known coding domain through ranked synonymous codon modifications. With the CORPSE method we demonstrated the 99 % reduction of *pB* activity within a reporter system without altering amino acid sequence of the overlapping gene. The CORPSE method was also applied to the intragenic promoter *trpCp* within *trpD* and eliminated 81 % of *trpCp* activity. We also inverted the CORPSE ranking system to create novel reverse promoters internal to the mCherry coding sequence. In one of our constructs, iCORPSE increased sfGFP reporter expression by > 10,000 %.

The CORPSE system shows promise in the future refactoring of complex genetic circuits by eliminating intragenic promoter sequences without potentially deleterious large codon usage changes which facilitates increased standardisation of parts. The CORPSE system has also demonstrated a future application in creating compressed and theoretically evolutionarily robust circuits through the embedding of a reverse promoter sequence internal to an existing coding sequence.

## Materials and Methods

### Calculating promoter frequencies

All σ^70^ promoter sequences present in the database were extracted from RegulonDB ^25^ (http://regulondb.ccg.unam.mx) via the σ70 sigmulon dataset. The sequence datasets were manually curated to remove non-hexameric promoter sequences. Non-hexameric sequences accounted for 8.5 % and 4.9 % of the −35 and −10 datasets, respectively. The −35 and −10 elements were extracted and used to calculate individual position specific scoring matrices (PSSM) ^45^ to generate log odds scores for each nucleotide in a position within the element, these log odds score were then reported as PSSM scores. The calculations were performed by curating −35 and −10 elements to only retain hexameric sequences. The nucleotides in each position in the sequence were counted and divided by the total number of sequences. These ratios were then divided by 0.25 to account for random frequency of an individual nucleotide occurring in the position. The normalised ratios had their binary logarithm values calculated to determine the log odds score of each nucleotide in the matrix. The frequency and PSSM (log odds) scores of each nucleotide at each position is presented in Supplementary File S1.

### Codon Restrained Promoter Silencing (CORPSE) algorithm

The CORPSE algorithm consisted of the following steps: (1) calculate log odd scores for hexameric −10 and −35 sequences, (2) separate sequences into codon triplets, (3) determine synonymous codon changes possible at each location, (4) combinatorically assemble all possible synonymous codon options, (5) calculate all log odds scores for each sequence, (6) output ranked sequences by log odd score.

Log odd scores used as a proxy for predicted promoter strength by assuming that a higher score contained more nucleotides required to facilitate the σ^70^-DNA interaction driving transcription. The CORPSE algorithm for *E. coli* is available as a webtool (https://bio-tools.com.au/promoter.html).

### Bacterial strains and plasmids

Three plasmid constructs were used throughout this study: pJ804 ^8^ used as the basis for CORPSE, pGERC ^31^ for the *trpCp* and iCORPSE studies, and pQE60 for the phenotypic analysis of *trpCp* modifications. All strains were grown in lysogeny broth (LB) and supplemented with either 100 μg/mL carbenicillin (pJ804), or 50 μg/mL kanamycin (pGERC) depending on which plasmid was required All plasmids were originally transformed and sequence verified in NEB Turbo (New England Biosciences, #C2984H). Plasmids were purified and transformed into the one of three bacterial strains if required. The NCTC122 *E. coli* strain, was used for all plasmids used in the CORPSE experimentation using TSS heat transformation ^46^. The pQE60::*trpCp* variants were transformed into the Δ*trpD* JW1255-1 strain (BW25113) from the KEIO collection ^47^ using the Mix & Go! protocol from Zymo Research (Zymo Research, #T3001).

### Designing CORPSE promoter constructs

Wild-type and CORPSE-designed promoter sequences were analysed in a reporter plasmid (pJ804) which drove sfGFP expression under BCD2 control ^44^ whilst mCherry under pCAT (BBa_I14033) constitutive expression with a consensus RBS sequence acted as an internal control. All promoter sequences from CORPSE analyse were embedded with 5` primer overhangs containing the variant promoter elements and native spacer sequence for assembly through round the horn (RTH) PCR ^44, 48, 49^ with a universal reverse phosphorylated primer. RTH variant primers were phosphorylated with NEB T4 Polynucleotide kits (New England Biosciences, #M0201) and 10 μM of RTH primers were used to amplify and mutate pJ804 backbone with Q5 Hot Start HF 2x Master Mix (New England Biosciences, #M0494). Template DNA was digested via DpnI (New England Biosciences, #R0176). Linear mutated DNA was ligated with T4 ligase (New England Biosciences, #M0202). Ligated plasmids were heat transformed into TSS competent cells ^46^ and recovered at 37 ^o^C for 1 h. All recovered cells were centrifuged at 6,000 RCF for 5 min (Eppendorf 5424), resuspended in 100 μL of media, and plated onto selective media.

### Designing *trpCp* promoter constructs

The *trpCp* sequence within *trpD* was analysed through CORPSE which generated two alternative promoter sequences. Only one of these sequences reduced the CORPSE score and was selected as the variant sequence for experimentation. Two gene constructs for *trpD* were synthesized (IDT), one containing the wild-type sequence (*trpCp-WT*) and one containing the variant sequence (*trpCp-V1*). The *trpD* sequences were inserted upstream of *sfGFP* in the pGERC plasmid immediately adjacent to the BCD2 RBS sequence, and immediately downstream of the T5 promoter in the pQE60 plasmid.

### Designing iCORPSE promoter constructs

The reverse complement sequence of mCherry was analysed through BPROM bacterial promoter prediction software ^30^ and two promoter sites were predicted (BPROM-1, and BPROM-2). These, along with two hexameric sequences separated by 17 nt and chosen for their alternative codons being similar to the σ^70^ consensus sequence (PROM-3), were analysed using the CORPSE method and the strongest promoter sequences were selected.

Using the pGERC plasmid ^31, 44^, separate DNA fragments were designed to remove the sfGFP EM7 promoter to silence sfGFP expression, and the four iCORPSE mutations were added (wild-type, V1, V2, and V3). These were assembled via NEBuilder HiFi DNA Assembly Master Mix (New England Biosciences, #E2621) and transformed into NEB Turbo competent cells (New England Biosciences, #C2984H), plasmid purified (QIAgen, #27104), and transformed into *E. coli* C strain NCTC122 TSS competent cells via the method described above.

### Flow Cytometry

Sequence confirmed strains were isolated as single colonies on LB agar plates (2 mM carbenicillin) and 1 mL cultures were grown overnight in a 96 well, square-welled, round bottomed deep well plate at 40 % liquid volume and sealed with a Breathe-easy sealing membrane (Merck, #2380059-1PAK). Overnight growth occurred in an Infors MT multitron pro shaker at 250 RPM rotating orbitally at 25 mm diametres at 37 °C for 17 h.

The density of 200 μL overnight cells were measured on a BMG Labtech SPECTROstar Nano plate reader with a CellStar clear 96 well plate. Cells were passaged to an OD600 of 1 in 1 mL and then diluted 1:100 into another deep well plate.

Cells were grown for 5 h to reach mid-log growth and the OD600 of 200 μL was measured on a BMG Labtech SPECTROstar Nano plate reader to confirm growth stage.

A further 200 μL was aliquoted into a CellStar clear 96 well plate and centrifuged in Eppendorf 5430R centrifuge with the plate swing bucket attachment at 2,240 RCF for 10 min to pellet cells. LB supernatant was removed and the cells resuspended in 1x phosphate buffered saline (PBS) media, and diluted 1:200 into 1x PBS media in a CellStar clear 96 well plate. Bacterial growth parameters have been reported to the best of our knowledge conforming with the MIEO v0.1.0 standard ^50^.

### Fluorescence Measurements

Fluorescence readings were performed as previously ^51^ on a Beckman Coulter CytoFLEX S (FITC, 488 nm excitation laser, 525/40 nm emission band-pass filter). 10,000 events were collected with acquisition settings of: FSC – 264, SSC – 2000, FITC – 299, and ECD – 150. Data was processed using CytExpert and data analysis conducted with median FITC and ECD scores and their generated standard deviations. FITC scores were normalized against ECD scores to create sfGFP/mCherry expression ratios, which were then divided against wild-type ratios to determine percentages of expression.

### RNA purification

RNA was purified from 20 mL *E. coli* cultures at early growth states (30 min post inoculation) using the RNeasy Mini kit (Qiagen: #74106) according to the manufacturer’s instructions, with the optional DNase on-column digestion step (Qiagen: #79254). Cultures were grown in an Infors MT multitron pro shaker at 250 RPM rotating orbitally at 25 mm diametres at 37 °C for both 17 h overnight step and for 30 min growth step.

### Quantitative PCR analysis

Purified RNA samples were reverse transcribed (RT) into cDNA according to the manufacturer’s instructions (ThermoFisher Scientific, #4368814). qPCR of the RT cDNA was performed according to the manufacturer’s instructions in a LightCycler 480 II (Roche Life Science using SYBR GREEN (Roche Life Science, #04707516001) with a 10 μL total reaction volume. sfGFP RNA expression was quantified using primers (F: GGTGAAGGTGACGCAACTAATGGTA, R: TTGGCCGACTCTGGTAACGACGCTG) normalized to the housekeeping gene cysG (F: GAAAGCCTTCTCGACACCTG, R: CGTTACAGAAGATGCGACGA) using the 2^−ΔΔC^T method ^52^.

### *TrpD* phenotype assay

Cultures of pQE60 containing strains (pQE60::*trpCp-WT*, pQE60::*trpCp-V1*, pQE60::*empty*) were grown overnight in an Infors MT multitron pro shaker at 250 RPM rotating orbitally at 25 mm diametres at 37 °C for 17 h in LB selective media. All cultures were normalized to an OD_600_ of 1 and serial diluted ten-fold until a final concentration of 1 × 10^−8^ was achieved. Dilutions were pipetted (2 μL aliquots) onto pre-warmed (37 °C) M9 agar plates. Plates were incubated overnight at 37 °C and assessed for strain growth.

## Acknowledgements

We recognize that the intellectual and physical labour of this research was conducted on the traditional lands of the Wattamattagal clan of the Darug nation and of the Gadigal and Wangal peoples of the Eora nation. We thank Thomas Williams for his suggestion to try inverting the CORPSE system to create new promoters via iCORPSE and Hannah Zhu for helpful suggestions to improve the manuscript.

