## Supplementary for "A Codon Constrained Method for Both Eliminating and Creating Intragenic Bacterial Promoters"

**CONTENTS**

SUPPORTING FIGURES

Figure S1: Predicted structures of transcription control elements within øX174 genome.

SUPPORTING TABLES

Table S1: Combinatorial designs for CORPSE *pB* promoter variants

SUPPORTING FILES

Supporting File S1: Position Specific Scoring Matrix data and Manual CORPSE analysis

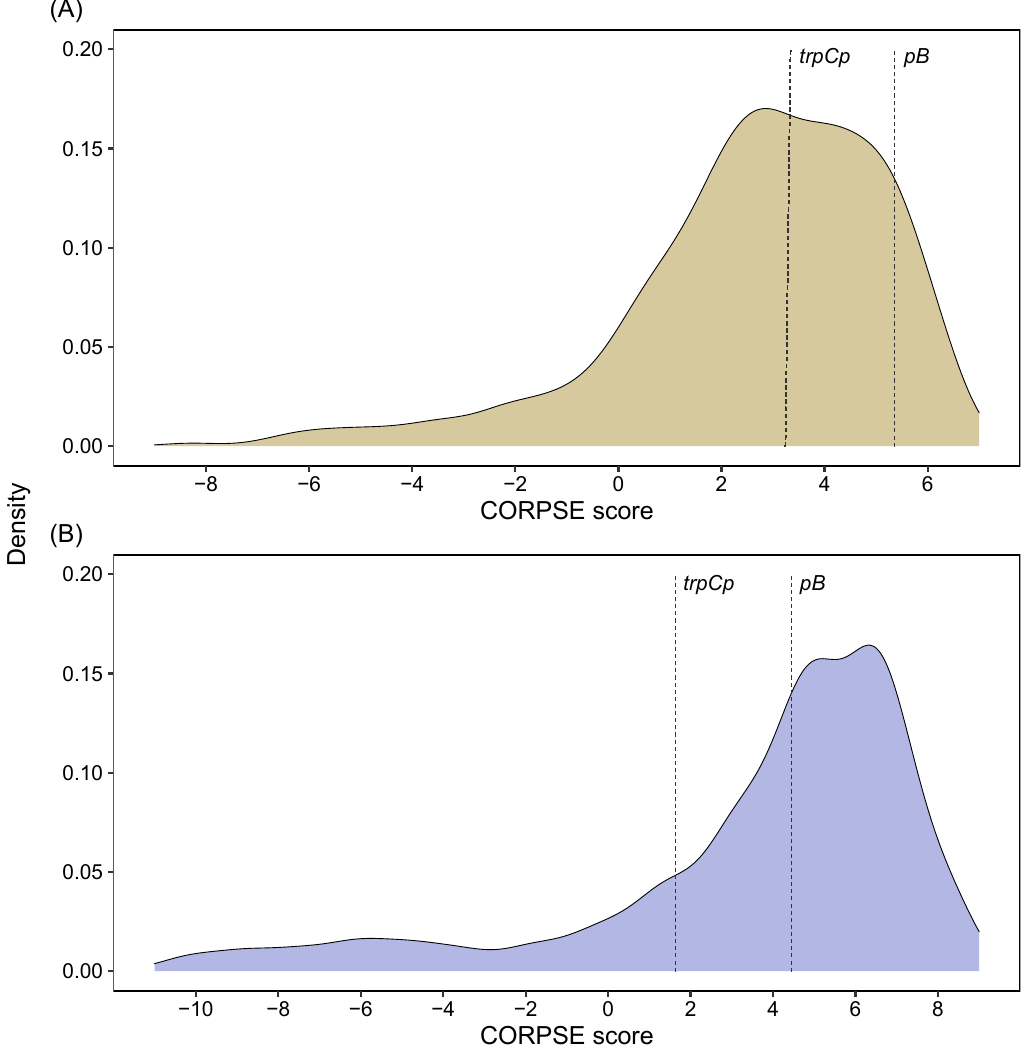

**Fig. S1. PSSM score distributions for all *E. coli* K12 promoter elements.** All promoter sequences were extracted from the curated RegulonDB dataset and used to calculate log odds scores of a position specific scoring matrix (PSSM). Curated dataset excluded non-hexameric sequences which accounted for 8.5 % and 4.9 % of the –35 and –10 datasets respectively. The calculated CORPSE scores derived from the PSSM for the wild-type *pB* and *trpCp* sequences are shown.

**Table S1.** Combinatorial designs for CORPSE *pB* promoter variants

| Promoter variant | –35 element | –10 element | PSSM score | | | | |
| --- | --- | --- | --- | --- | --- | --- | --- |
|  |  |  | –35 | Percentile | –10 | Percentile | combined |
| CORPSE-WT | TTGCAA | TACAGT | 5.4 | 89^th^ | 4.5 | 44^th^ | 9.9 |
| CORPSE-01 | TTGCAA | TACTCC | 5.4 | 89^th^ | -0.2 | 15^th^ | 5.2 |
| CORPSE-02 | TTGCAA | TACTCG | 5.4 | 89^th^ | -0.7 | 13^th^ | 4.7 |
| CORPSE-03 | CTACAA | TACTCC | 0.9 | 20^th^ | -0.2 | 15^th^ | 0.7 |
| CORPSE-04 | CTACAA | TACTCG | 0.9 | 20^th^ | -0.7 | 13^th^ | 0.2 |
| CORPSE-05 | CTACAG | TACTCC | -0.1 | 12^th^ | -0.2 | 15^th^ | -0.3 |
| CORPSE-06 | CTACAG | TACTCG | -0.1 | 12^th^ | -0.2 | 15^th^ | -0.3 |
| CORPSE-07 | CTCCAA | TACTCC | 1.0 | 20^th^ | -0.2 | 15^th^ | 0.8 |
| CORPSE-08 | CTCCAA | TACTCG | 1.0 | 20^th^ | -0.7 | 13^th^ | 0.3 |
| CORPSE-09 | CTCCAG | TACTCC | 0.0 | 12^th^ | -0.7 | 13^th^ | -0.7 |
| CORPSE-10 | CTCCAG | TACTCG | 0.0 | 12^th^ | -0.7 | 13^th^ | -0.7 |
| CORPSE-11 | CTGCAG | TACTCC | 1.7 | 28^th^ | -0.2 | 15^th^ | 1.5 |
| CORPSE-12 | CTGCAG | TACTCG | 1.7 | 28^th^ | -0.2 | 15^th^ | 1.5 |
| CORPSE-13 | CTTCAA | TACTCC | 0.6 | 17^th^ | -0.2 | 15^th^ | 0.4 |
| CORPSE-14 | CTTCAA | TACTCG | 0.6 | 17^th^ | -0.7 | 13^th^ | -0.1 |
| CORPSE-15 | CTTCAG | TACTCC | -0.4 | 11^th^ | -0.2 | 15^th^ | -0.6 |
| CORPSE-16 | CTTCAG | TACTCG | -0.4 | 11^th^ | -0.7 | 13^th^ | -1.1 |
| CORPSE-NC | GCAGGG | ACGCTC | -11.8 | N/A | -12.4 | N/A | -24.2 |

All promoters used identical spacer sequences (aatacgtggccttataggt). The percentile value for both the –35 and –10 consensus sequences are 99^th^ percentile. N/A represents values below the 1^st^ percentile.
